# Early-evening biting by *Anopheles stephensi* in Southern Ethiopia: A Challenge for Bed Net Use as Vector Control in Africa

**DOI:** 10.1101/2025.08.25.672230

**Authors:** Maxwell G. Machani, Samuel S. C. Rund, Dawit Hawaria, Guiyun Yan

## Abstract

*Anopheles stephensi* is rapidly expanding across Africa, posing new challenges for malaria control. Its biting time patterns, however, remain poorly characterized, raising uncertainty about the effectiveness of bed nets against this invasive vector. To address this gap, we investigated diel biting activity, feeding propensity, and flight behavior using complementary behavioral assays on females reared from wild-caught larvae in Hawassa City, southern Ethiopia. Flight monitoring assays revealed that *An. stephensi* exhibited pronounced activity at dusk, beginning between 18:00 and 19:00 with the onset of scotophase, and little to no activity during the photophase. Blood-feeding propensity, defined as the proportion of mosquitoes taking a blood meal when offered, peaked during the early scotophase (18:00-22:00) at 33.3-51.7%, but was markedly reduced during daylight hours (0-16.7%). Human landing catches in large-cage enclosures confirmed this early evening activity: 83.5% of total landings occurred between 18:00 and 22:00 with a sharp peak at 18:00-19:00, corresponding to a mean biting rate of 17.8 bites per person per hour. These findings demonstrate that invasive *An. stephensi* primarily seek hosts and bite during the early evening, a time when people are often unprotected by bed nets. This behavior reduces the protective impact of conventional net-based interventions and underscores the need for African National Malaria Control Programs to deploy complementary measures such as spatial repellents and larval source management to mitigate early-evening transmission. Moreover, this study highlights the utility of integrated behavioral assays for estimating biting time, offering approaches that can be extended to other vector species across Africa.

## Introduction

*Anopheles stephensi*, traditionally confined to Asia and the Middle East, has recently invaded multiple African countries, raising significant concerns for urban malaria transmission (1–4). Geospatial modeling suggest that its continued spread could expose over 126 million people to malaria risk in African urban settings (5), potentially reversing progress made in malaria control in sub-Saharan Africa. In Ethiopia, models predict a 50% rise in *Plasmodium falciparum* malaria cases without scaling up of interventions like long-lasting insecticidal nets (LLIN), indoor residual spraying (IRS) coverage and larval source management (LSM) (6). Although some countries have initiated responses to the invasion, critical knowledge gaps remain. Most notably, there is limited understanding of *An. stephensi* feeding behavior in African settings, as field observations of adult mosquitoes are scarce, constraining the development of effective, evidence-based control strategies (7,8).

Mosquito activity is regulated by an internal circadian clock, which interacts with environmental signals such as light, temperature, and host availability to control critical behaviors like host-seeking and biting (9,10). These activity rhythms have direct implications for vector control effectiveness. For instance, bed nets are predicated on the assumption that vectors bite indoors during the late-night hours when most people are asleep (11). For decades, this assumption was largely valid, as the primary African malaria vectors, *Anopheles gambiae* and *An. funestus*, exhibited peak biting activity from midnight through the late-night hours. This alignment maximized human protection under bed nets while also ensuring high mosquito mortality through contact with insecticide-treated surfaces (12,13). However, ecological and evolutionary shifts in mosquito biting time or host-seeking patterns can undermine the effectiveness of current control tools. Altered biting rhythms in African vectors alone are estimated to account for an additional 10.6 million malaria cases annually (14,15). These behavioral changes may arise from intrinsic biological factors or evolve under selective pressure from widespread LLIN deployment, enabling mosquitoes to exploit times or locations where people are less protected (16).

Following the invasion of *An. stephensi* into several African countries, National Malaria Control Programs (NMCPs) have expanded surveillance efforts and implemented additional interventions targeting both adult and larval stages in affected areas (17). Despite these efforts, most knowledge about *An. stephensi* behavior derives from studies conducted decades ago in its native Asian range where biting patterns are often multimodal or arrhythmic (18,19). Limited understanding of biting and host-seeking behaviors of invasive *An. stephensi* populations in Africa creates uncertainty about the effectiveness of LLINs in mitigating its impact on malaria transmission. Here, we investigate the temporal biting patterns (the timing of foraging decisions in a large enclosure with host cues present), biting propensity (the proportion of mosquitoes initiating feeding when exposed to a short-range host stimulus), and flight activity (a proxy for host-seeking behavior in the absence of host cues) of field-collected *An. stephensi* from Ethiopia. Using three complementary approaches in both laboratory and semi-field (large enclosure) settings, this work provides evidence to guide African NMCPs in assessing the potential effectiveness of bed net use as a control measure in regions affected by this invasive vector.

## Methodology

### Study site and mosquito collection

*Anopheles* larval collections were conducted from March to July 2025 in Hawassa City (7.058873°N, 38.48982°E, altitude 1,708 meters (5,604 feet) above sea level) located in southern Ethiopia along the shores of Lake Hawassa. The area experiences a short rainy season from March to May, followed by a longer wet season from July to October. Malaria transmission in the area is seasonal, with peak transmission typically occurring between September and December (8). *Anopheles* larvae were collected from a variety of urban breeding habitats in Hawassa, including unmaintained swimming pools, brick-making pits, water storage containers, and tire tracks. They were reared under standardized laboratory conditions at Hawassa University insectary, maintained at 27 ± 2 °C, 75 ± 5% relative humidity, and a 12:12 h light:dark cycle. Larvae were fed daily on finely ground Tetramin® fish food and raised in rainwater. Emerging adults were maintained on 10% sucrose solution. Upon emergence, *Anopheles* mosquitoes were morphologically identified using the update anopheline keys (20). Only *An. stephensi* identified morphologically were used for subsequent assays.

### Biting propensity

This assay measures the mosquitoes’ short-range biting drive by assessing what proportion blood feed, when presented with a host (*e.g.,* a human arm) in close proximity for a fixed time period. Thirty mated, 6–8-day-old, non-blood-fed *Anopheles stephensi* females (F_0_ generation) were aspirated into seventeen paper cups, provided with 10% sucrose, and allowed to acclimate overnight in an unoccupied room in the field. Starting at 6:00 PM the following day, blood-feeding was assessed every 3 hours for 48 hours by exposing a randomly selected cup (30 mosquitoes) to the investigator’s arm for 6 minutes, using the method of Rund et al. (21). After each exposure, mosquitoes were frozen and visually examined for the presence of any blood in the abdomen. The experiment was conducted under ambient fluctuating temperature and humidity with a 12:12 light:dark cycle provided through the windows from the natural environment. No artificial lighting was used at any time during the 48-hour period, allowing the room to reflect natural lighting conditions typical of rural homes, bright during the day through natural openings and dark after sunset.

### Biting rhythms in large enclosure

This assay allowed us to better approximate mosquitoes’ host-seeking drive by enabling them to express a range of behaviors (they have to actively move toward the host within the enclosure and attempt to bite). We conducted human landing collections (HLC) within a portable semi-field enclosure that simulates natural host-seeking conditions while maintaining containment of this invasive species (Figure S1.A). This approach addressed two key limitations: (i) the low densities of *An. stephensi* currently observed in the field, and (ii) ethical and biosafety concerns associated with releasing non-native vectors into open environments. The experimental system consisted of a pop-up tent (3.66 x 3.66 M floor dimension, 2.4 M peak height, East Oak) enclosure constructed using netting to allow airflow and natural ambient light while ensuring complete containment. Female mosquitoes aged 6-8 days (F_0_ generation), non-blood-fed, and sugar-starved for two hours (with access to moist cotton) were used for the assays. Each evening, 100 adult females were released into the enclosure at 17:00 h, one hour prior to the start of collections, to allow acclimatization. HLCs were conducted from 18:00 h to 08:00 h across two consecutive nights. A single human volunteer, seated at the middle of the enclosure, collected landing mosquitoes using a mouth aspirator. Fifteen-minute breaks were provided at the end of each hour to minimize collector fatigue and maintain consistency.

### Flight activity monitoring

The onset and duration of nocturnal flight activity (a proxy for the host-seeking flight behavior) of wild *An. stephensi* was monitored using a Locomotor Activity Monitor 25 (LAM 25) system (TriKinetics, Waltham, MA) (Figure S1B) as previously described by Rund et al (22). Briefly, non-blood-fed 6-8 days old (F_0_ generation), Individual mosquitoes were placed in 25 x 150 mm clear glass tubes with access to 10% sucrose in the tubes provided *ad libitum*. Flight activity was recorded as infrared beam breaks per minute on a per-animal and per-minute basis. Each LAM unit allows simultaneous monitoring of 32 mosquitoes in a 4 x 8 vertical by horizontal matrix. All recordings occurred in a light-proof room with its own lighting system in a 12 hr:2 hr light:dark cycle. Light was provided by a white LED light bar.

### Data analysis

Locomotor flight activity was visualized in one-minute bins using ClockLab version 6 (Actimetrics, Wilmette, IL) and behavioral patterns were presented as actograms. Animals were considered alive and healthy for the duration of the analysis if they displayed at least 25 beam/breaks per day and the last day of activity had no less than 20% of the average daily beam/breaks per day for that animal. Human landing rates were used to estimate human biting rates by dividing the total mosquitoes collected by the number of persons per night. Nocturnal biting activity was expressed as the mean number of landings per person per hour, assuming hourly catches represent mosquitoes attempting to feed during that period. Blood-feeding propensity was calculated as the proportion of mosquitoes with visible blood meals in the gut relative to the number exposed at that time.

## Results

### Flight behavior rhythms

To better understand the endogenous flight activity rhythms in the absence of host cues, we monitored the flight/locomotor activity rhythm of individual mosquitoes on a minute-by-basis basis using a Locomotor Activity Monitoring (LAM). Under diel light:dark (LD) conditions, *An. stephensi* flight activity exhibited a robust daily rhythm, with activity onset at dusk between 18:00 and 19:00 marking the beginning of the scotophase (dark period of a light:dark cycle) (Figure 1). Activity peaked early in the scotophase and gradually declined toward midnight, remaining low throughout the photophase (light period of a light:dark cycle). This circadian-regulated pattern was consistently maintained over the five-day monitoring period (Figure 1).

**Figure 1:**
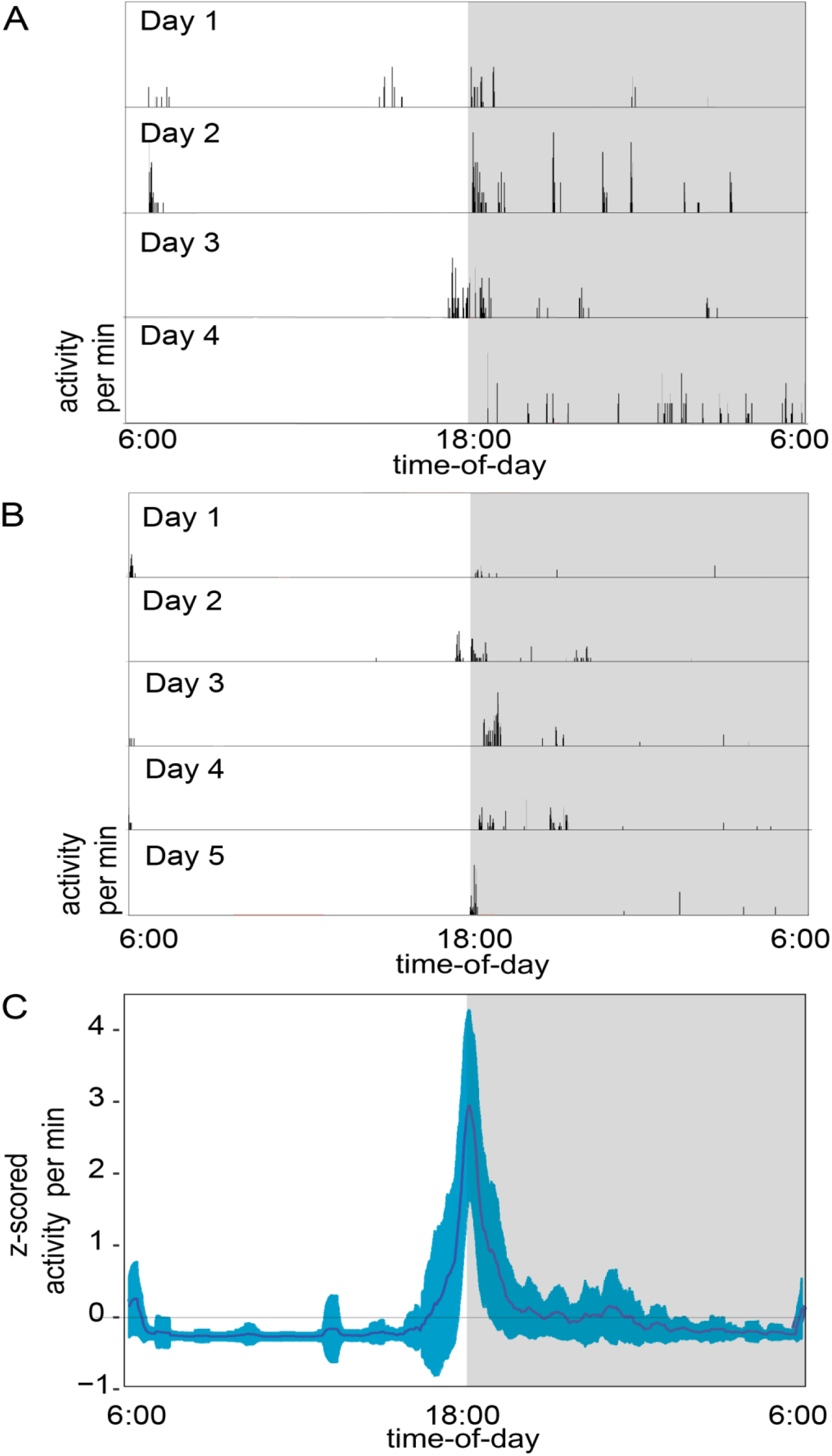
Flight activity recording in locomotor activity monitors (LAM25) demonstrates *An. stephensi* is most active at dusk. Mosquitoes were monitored over several days in two separate runs. The grey/white backgrounds indicate the 24h light:dark photoperiod. A and B are two separate runs and are representative actiograms of a single mosquito monitored continuously. C is a z-scored average of activity of the second run that collapses 5 days of activity from the 26 surviving animals into a single representative temporal profile. The blue is the standard deviation.

### Blood-feeding propensity rhythms

Blood feeding rhythms also varied significantly across the diel cycle over the 48-hour period (Figure 2). The highest feeding rates were observed during the early evening hours (18:00-22:00), with blood feeding proportions ranging from 33.3% (95% CI: 15.5-51.1) to 51.7% (95% CI: 33.5-69.9) of the mosquitoes biting that were offered a blood-meal at short range. Feeding rates declined during the late night to early morning hours (00:00-06:00), with proportions between 17.9% (95% CI: 3.8-32.0) and 30.4% (95% CI: 11.6-49.2). The lowest feeding activity (17.9%) was recorded in the late morning, while daylight hours (09:00-15:00) consistently showed minimal feeding, ranging from 0% to 16.7% (95% CI: 3.3-30). Notably, the peak blood-feeding period closely coincided with the onset of flight activity. suggesting a coupling of the behaviors monitored.

**Figure 2:**
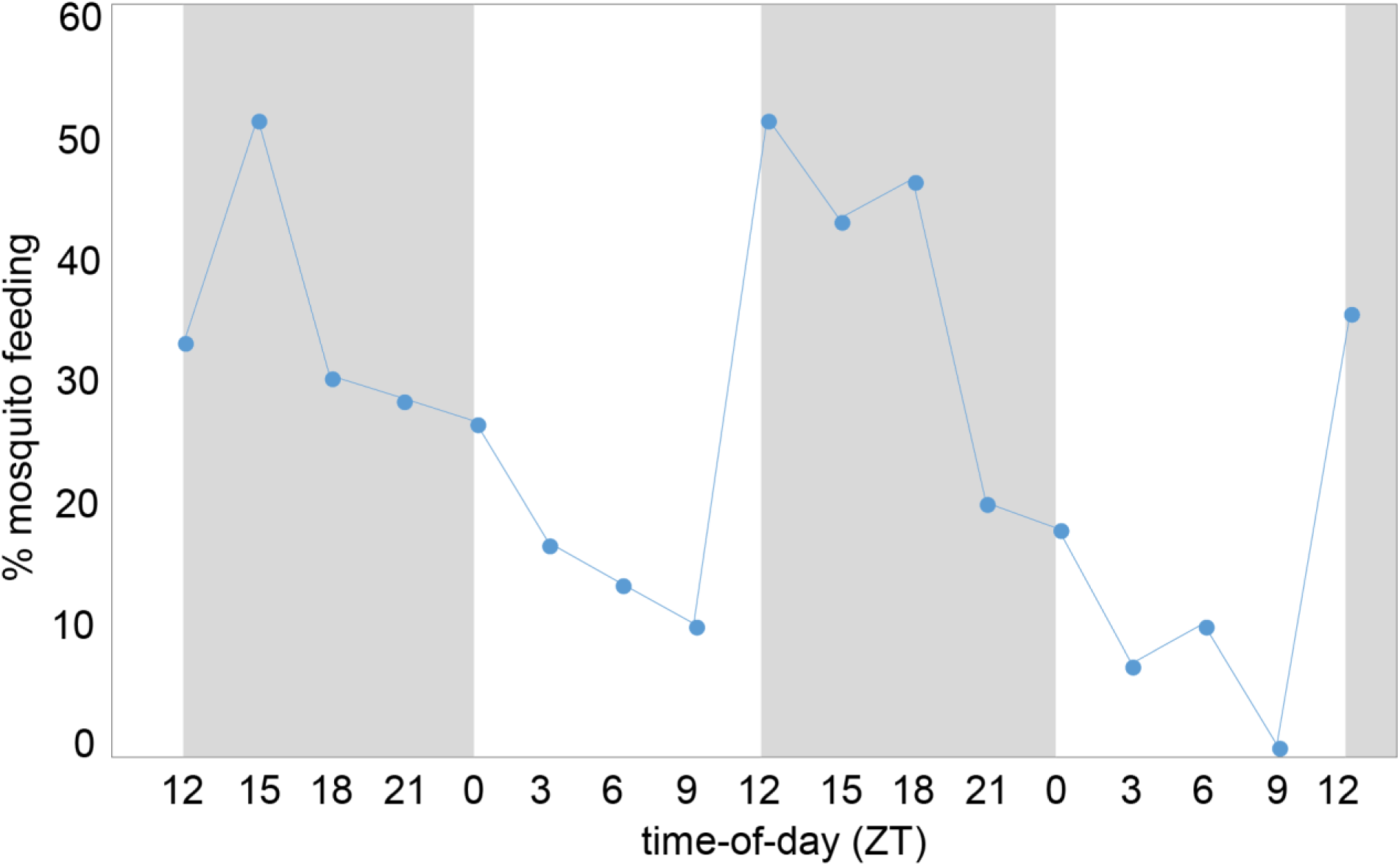
Blood-feeding propensity of *An. stephensi* mosquitoes. Every 3h a separate pot of 30 female mosquitoes was exposed to a human arm. The proportion of mosquitoes that blood-fed during each 6-minute exposure to a human arm under normal light-dark (LD) conditions is indicated. The grey/white backgrounds indicate the 24h light:dark photoperiod.

### Host seeking activity

Human landing catch (HLC) data collected in a large enclosure revealed a sharply skewed temporal biting pattern of *An. stephensi*, characterized by a pronounced early evening peak (Figure 2). Overall, *An. stephensi* exhibited intense host-seeking activity during the early evening hours, with 83.5% (95% CI: 77.7-89.5) of total mosquito landings recorded between 18:00 and 22:00 h with a pronounced and distinct peak between 18:00 and 19: 00 h (mean 17.8 bites/person/hour). Biting activity declined steadily from 23:00 to 04:00 h, with landing rates ranging from 0.63% (0.25 mean bites/person/hour) to 5.14% (2 mean bites/person/hour). Minimal to no activity was recorded between 05:00 and 07:00 h. The early-biting window observed here was consistent with blood-feeding propensity and the onset of flight activity.

## Discussion

The successful invasion of disease vector species depends on its capacity to establish and expand within novel environments. This success is shaped by a combination of intrinsic biological traits, such as host-seeking behavior, and extrinsic factors, including host availability, climatic conditions, and human-driven environmental changes (23,24). Understanding the bionomics of these species is essential for designing effective vector surveillance, avoidance and control strategies (25). For instance, detailed knowledge of biting behavior can inform the timing and selection of personal protection measures aimed at reducing human-vector contact. In this study, we show through three complementary approaches in both laboratory and large enclosure settings that wild *An. stephensi*, an invasive vector of Asian origin, exhibits early evening biting and host-seeking activity in Ethiopia. This behavior underscores a potential limitation of LLINs, which are designed to primarily protect against late-night indoor biting.

Mosquito behaviors such as host seeking, blood feeding, mating, and oviposition are governed by circadian rhythms (21,26,27), which are regulated by an endogenous circadian clock (21,27,28) that synchronizes key activities with optimal environmental conditions. Our three complementary host seeking/blood feeding assays showed strong alignment in results. We observed that the blood-feeding activity of *An. stephensi* occurred primarily during early evening hours (18:00-22:00 h), with peak activity between 18:00 and 19:00 h. This feeding rhythm remained consistent over multiple observation periods. A similar pattern was observed in a large enclosure setup, where most mosquito landings on a human host occurred during the first half of the night, again peaking at 18:00-19:00 h and declining steadily thereafter (Fig. 3). Finally, using the Locomotor Activity Monitoring (LAM) system, we demonstrate that wild *An. stephensi* flight activity is most active at dusk, with occasional dawn activity over the five-day monitoring period.

**Figure 3:**
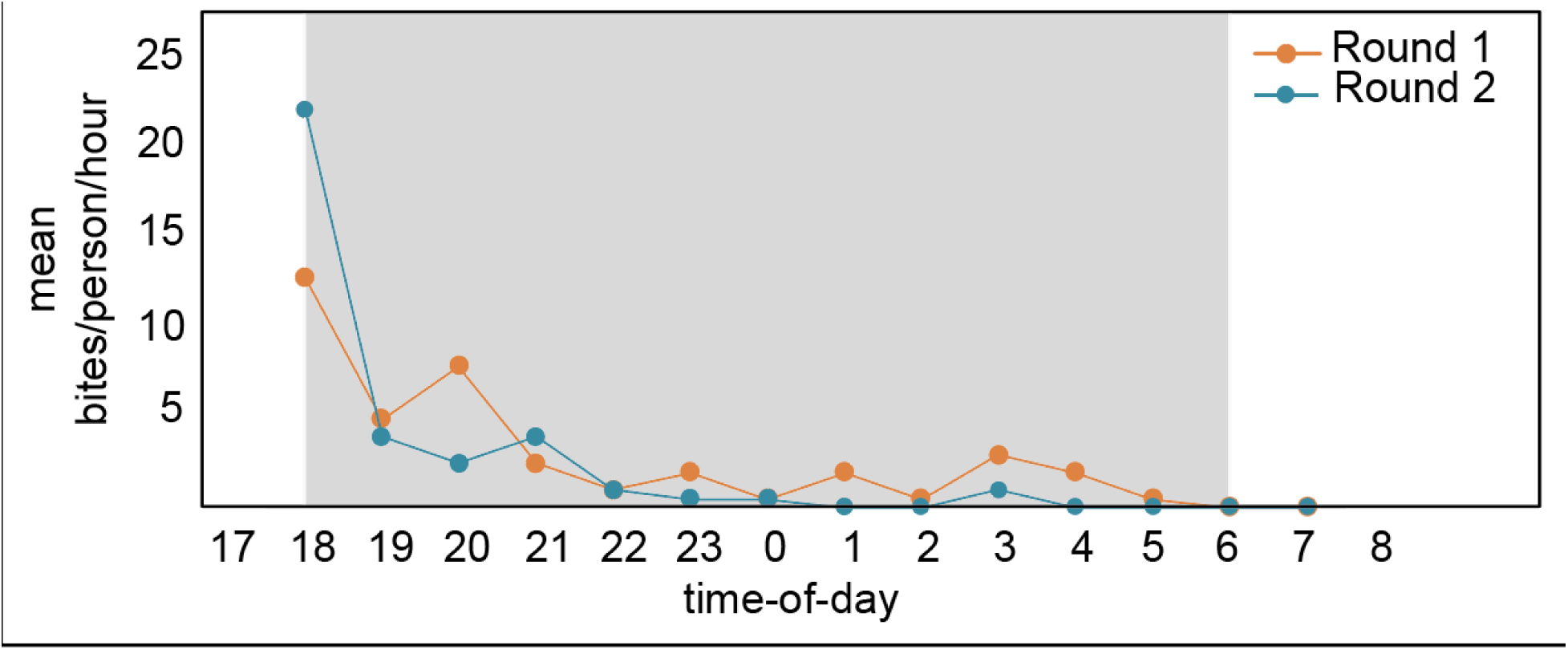
Host-seeking and biting rhythms. Hourly human landing rates of *An. stephensi* released in a large enclosure from 18:00 to 08:00 under natural light-dark conditions. Values represent the mean number of mosquitoes landing per person per hour. The grey/white backgrounds indicate the 24h light:dark photoperiod.

These findings indicate a highly concentrated biting window during the early evening hours which suggest that Ethiopian *An. stephensi* will preferentially bite during early evening hours, coinciding with periods when people are active and unprotected by LLINs. Studies in Ethiopia have observed that about 83% of the population go to sleeps after 21:00 h (29,30). Our findings contrast with earlier reports from *An. stephensi’s* native range which suggests plasticity in its biting behavior, varying by location and environmental conditions. Only a limited number of studies have documented these variations and some are decades ago. For example, in India, *An. stephensi* has been reported to exhibit peak biting during the early morning hours in the pre-monsoon period, while during the monsoon season, two distinct biting peaks were observed from 20:00 to 00:00 h and from 03:00 to 06:00 h (18). In southern Iran, *An. stephensi* was observed biting throughout the night, with a peak around midnight (00:00 h) (19). Similarly, this present work contrasts with our previous study on laboratory colonies of *An. stephensi* (originally from Pakistan) with nocturnal flight activity starting at the mid-night, and not at dusk (9). These variability in results may also suggest local adaptation of *An. stephensi* in Ethiopia, with the mosquito adjusting its biting behavior to differing environmental factors and ecological pressures in its invasive range (31). Differences from previous reports may also be attributed to our use of F_0_ mosquitoes raised from wild-collected larvae in Africa, which retain natural circadian rhythms better than long-established laboratory colonies prone to genetic bottlenecks and behavioral shifts (32,33). Interestingly, no early morning biting was observed for this vector, a major concern for the native vectors (34–36) highlighting how biting patterns and risks differ across vector species. Our findings underscore the importance of understanding behavioral differences and changes to develop effective, region-specific control strategies.

The onset and increase in flight activity recorded using the locomotor activity monitors, correspond with increased blood feeding propensity and biting rates observed in parallel assays, confirming the early-biting behavior of *An. stephensi* in southern Ethiopia. Moreover, this synchronization between flight activity and host-seeking behavior suggests that locomotor activity monitoring can serve as a reliable, cost-effective, and more standardized proxy for estimating biting risk in its invasive range. Unlike HLC which is often associated with ethical concerns, this approach provides a practical and feasible alternative, especially in difficult surveillance contexts, and supports the development of targeted vector control strategies.

The findings of this study underscore a limitation of current vector control strategies that rely heavily on long-lasting insecticidal nets (LLINs), which primarily protect individuals during sleeping hours (37). While LLINs remain effective as both a mechanical and insecticidal barrier during sleep, their utility is limited when significant transmission occurs before bedtime (38,39). Our results suggest that significant vector-host contact and hence potential transmission occurs in the early evening before individuals typically go to bed. This early-biting behavior aligns closely with active human routines, increasing exposure risk and diminishing the protective window offered by LLINs. Although we did not assess human behavioral patterns in this study, a recent report from Ethiopia found that 85% of people were outdoors or indoors but awake between 18:00 and 22:00, overlapping with the peak biting times we observed for *An. stephensi* (30). Moreover, *An. stephensi* has been shown in Ethiopia to transmit both *Plasmodium falciparum* and *P. vivax* in urban settings (40,41) underscoring its potential to drive urban malaria unlike native species *An. arabiensis*, which show similar behavioral plasticity (29) but less capable of exploiting urban habitats (42). To bridge this protection gap, personal protection tools, *e.g.* LSM (larval source management), mosquito repellents (both spatial and topical) and long-sleeved clothing, may offer protection during evening hours. Since people are likely to be outdoors during these peak biting times, as observed in Ethiopia (29,30), outdoor biting by this vector is highly likely. Spatial repellents, in particular, have demonstrated effectiveness in reducing malaria transmission in endemic areas (43). This calls for a strategic shift: rather than merely intensifying LLIN distribution, NMCPs should integrate complementary tools tailored to the local biting behavior of this vector.

Our findings are based on data from semi-field experiments, as field studies have been constrained by the inability to consistently collect large number of *An. stephensi* due to its recent invasion and the ineffectiveness of current mosquito trapping methods. Standard surveillance tools (i.e., CDC light traps, BG-Sentinel, and human landing catches-gold standard) have not reliably captured host-seeking *An. stephensi* in either African settings (7,8) or its native range (18). To overcome these limitations, we used F_0_ adults reared from field-collected larvae and conducted experiments in laboratory and semi-field systems, which allowed for behavioral observations under near-natural conditions while controlling for confounding environmental variables (44,45). Given the current gaps in entomological data, our semi-field approaches can offer timely and ecologically relevant insights to guide response strategies. Nonetheless, we recommend longitudinal field studies with larger sample sizes across the invasive range to capture potential regional differences (*i.e.* local adaption and/or phenotypic plasticity) in early and late-biting patterns, coupled with human behavioral observations, to better understand the role of early-biting by *An. stephensi* in sustaining malaria transmission and to assess the impact of vector control tools.

## Conclusion

Through three complementary approaches, this study demonstrates that invasive *An. stephensi* populations in Ethiopia display pronounced early evening biting activity, with significant host-seeking occurring before people typically go to bed. This early-biting behavior increases the likelihood of outdoor exposure and diminishes the protective effectiveness of LLINs, leaving a window of vulnerability when individuals are unprotected. These findings underscore the importance for African National Malaria Control Programs to closely monitor *An. stephensi* biting patterns and to implement integrated strategies that supplement LLINs such as targeted spatial repellents, larval source management, and interventions that address outdoor biting, to mitigate early evening transmission risks. Moreover, the behavioral assays applied here offer valuable tools for assessing and tracking biting behaviors of other disease vector species, especially where standard human landing catch (HLC) methods are ineffective, thereby strengthening surveillance and enabling adaptive vector control across diverse ecological settings.

## Declarations

### Ethics approval and consent to participate

Ethical clearance was obtained from the Institutional Review Board (IRB) of College of Medicine and Health Science, Hawassa University and the University of California, Irvine Institutional Review Board (UCI IRB). Volunteer collecting mosquitoes in the large enclosure consented and was trained to collect mosquitoes as they land before biting. The mosquitoes used were raised from larvae with no risk of infection. For mosquito collection, oral consent was obtained from landowners in each location. These locations were not protected land, and the field studies did not involve endangered or protected species.

### Consent for publication

Not applicable.

### Competing interests

The authors declare that they have no competing interests.

### Availability of data and materials

The dataset supporting the conclusions of this article is included within the article.

### Funding

This work is funded by National Institute for Health (D43 TW001505 and U19 AI129326) to G.Y. Research equipment and travel funds to S.S.C.R. were funded by an Initiation Grant from the University of Notre Dame, Notre Dame Research office. S.S.C.R. is funded by the Center for Research Computing, University of Notre Dame. The funder had no role in study design, data collection and analysis, decision to publish, or preparation of the manuscript.

### Authors’ contributions

Conceptualisation: MGM, SSCR, G.Y. Methodology: MGM, SSCR, Investigation: MGM, SSCR, DH, Analysis: SSCR, MGM. Writing-original draft: MGM, SSCR. Writing-review and editing: MGM, SSCR, DH, G.Y. All authors read and approved the final manuscript.

## Acknowledgements

We thank the ICEMR Ethiopia technical staff, specifically Bereket Tiruye, Shibeshi Getachew, Eyasu Tekle and Sisay Mesele, for their assistance with field sample collection, and Hawassa University, for kindly letting us use their facilities.

---

**Figure S1:**
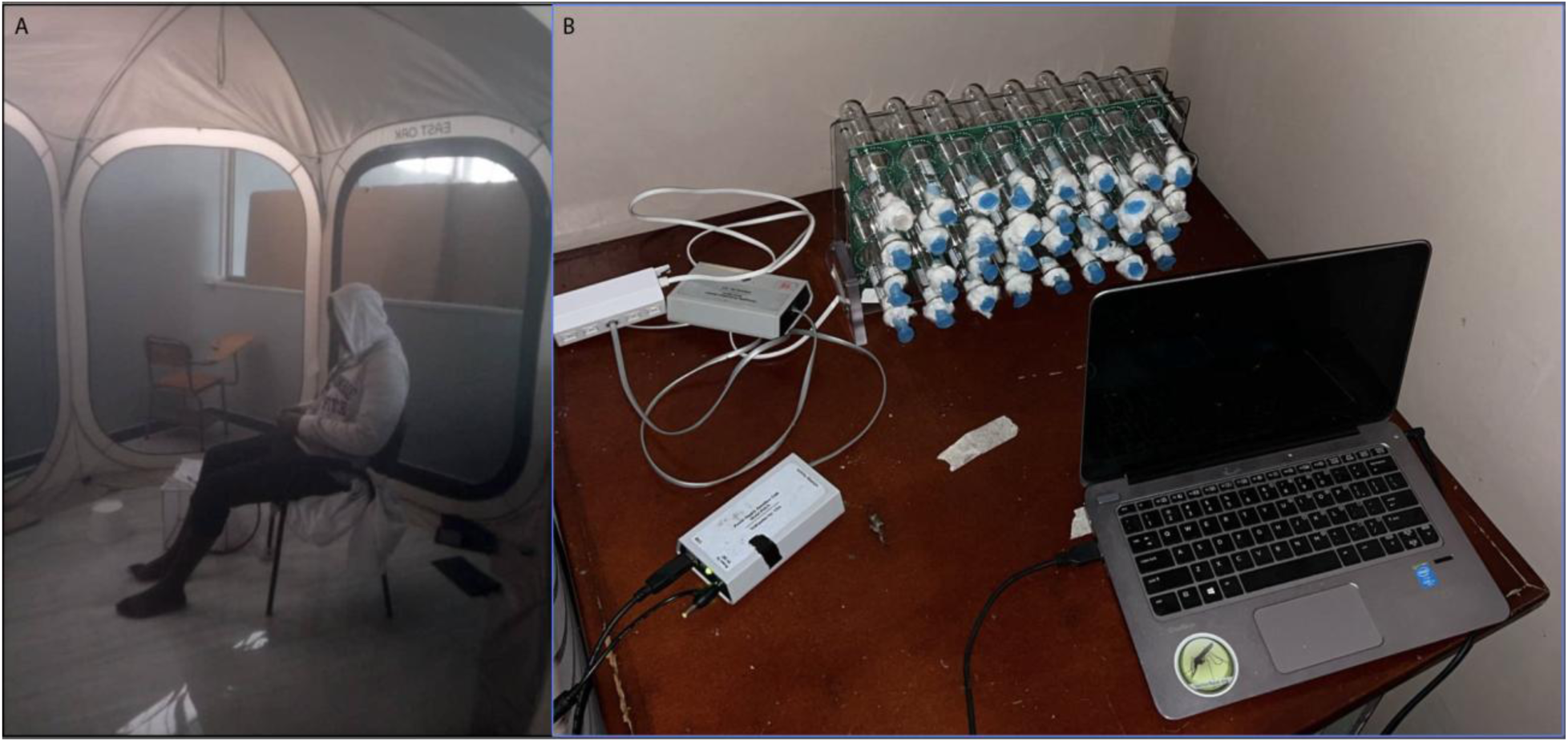
Experimental assays A: Human landing catches in a large enclosure-pop up tent (Mosquitoes in free range decide when to bite when a human host is present), B. Locomotor activity monitoring setup-used to continuously measure flight activity as a proxy for host-seeking behavior

## Notes

### Competing Interest Statement

The authors have declared no competing interest.

## References

1. Carter TE, Yared S, Gebresilassie A, Bonnell V, Damodaran L, Lopez K, et al. First detection of Anopheles stephensi Liston, 1901 (Diptera: culicidae) in Ethiopia using molecular and morphological approaches. Acta Trop. 2018 Dec;188:180–6.

2. Ochomo EO, Milanoi S, Abong’o B, Onyango B, Muchoki M, Omoke D, et al. Detection of Anopheles stephensi Mosquitoes by Molecular Surveillance, Kenya. Emerg Infect Dis. 2023 Dec;29(12):2498–508.

3. Faulde MK, Rueda LM, Khaireh BA. First record of the Asian malaria vector Anopheles stephensi and its possible role in the resurgence of malaria in Djibouti, Horn of Africa. Acta Trop. 2014 Nov;139:39–43.

4. WHO. Vector alert: Anopheles stephensi invasion and spread in Africa and Sri Lanka [Internet]. 2022 [cited 2025 Aug 11]. Available from: https://www.who.int/publications/i/item/9789240067714

5. Sinka ME, Pironon S, Massey NC, Longbottom J, Hemingway J, Moyes CL, et al. A new malaria vector in Africa: Predicting the expansion range of Anopheles stephensi and identifying the urban populations at risk. Proc Natl Acad Sci U S A. 2020 Oct 6;117(40):24900–8.

6. Hamlet A, Dengela D, Tongren JE, Tadesse FG, Bousema T, Sinka M, et al. The potential impact of Anopheles stephensi establishment on the transmission of Plasmodium falciparum in Ethiopia and prospective control measures. BMC Med. 2022 Apr 20;20(1):135.

7. Samake JN, Athinya DK, Milanoi S, Ramaita E, Muchoki M, Omondi S, et al. Spatial distribution and population structure of the invasive Anopheles stephensi in Kenya from 2022 to 2024. Sci Rep. 2025 June 6;15(1):19878.

8. Hawaria D, Kibret S, Zhong D, Lee MC, Lelisa K, Bekele B, et al. First report of Anopheles stephensi from southern Ethiopia. Malar J. 2023 Dec 8;22(1):373.

9. Rund SSC, O’Donnell AJ, Prior KF, van der Veen DR. Seasonal plasticity in daily timing of flight activity in Anopheles stephensi is driven by temperature modulation of dawn entrainment. Philos Trans R Soc B Biol Sci. 2025 Jan 23;380(1918):20230343.

10. Rund SSC, O’Donnell AJ, Gentile JE, Reece SE. Daily Rhythms in Mosquitoes and Their Consequences for Malaria Transmission. Insects. 2016 Apr 14;7(2):14.

11. Pates H, Curtis C. Mosquito behavior and vector control. Annu Rev Entomol. 2005;50:53–70.

12. Fontenille D, Lochouarn L, Diatta M, Sokhna C, Dia I, Diagne N, et al. Four years’ entomological study of the transmission of seasonal malaria in Senegal and the bionomics of Anopheles gambiae and A. arabiensis. Trans R Soc Trop Med Hyg. 1997;91(6):647–52.

13. Githeko AK, Adungo NI, Karanja DM, Hawley WA, Vulule JM, Seroney IK, et al. Some observations on the biting behavior of Anopheles gambiae s.s., Anopheles arabiensis, and Anopheles funestus and their implications for malaria control. Exp Parasitol. 1996 Apr;82(3):306–15.

14. Sanou A, Nelli L, Guelbéogo WM, Cissé F, Tapsoba M, Ouédraogo P, et al. Insecticide resistance and behavioural adaptation as a response to long-lasting insecticidal net deployment in malaria vectors in the Cascades region of Burkina Faso. Sci Rep. 2021 Sept 2;11(1):17569.

15. Sherrard-Smith E, Skarp JE, Beale AD, Fornadel C, Norris LC, Moore SJ, et al. Mosquito feeding behavior and how it influences residual malaria transmission across Africa. Proc Natl Acad Sci. 2019 July 23;116(30):15086–95.

16. Gatton ML, Chitnis N, Churcher T, Donnelly MJ, Ghani AC, Godfray HCJ, et al. The Importance of Mosquito Behavioural Adaptations to Malaria Control in Africa. Evolution. 2013;67(4):1218–30.

17. WHO. Partners convening: a regional response to the invasion of Anopheles stephensi in Africa. Meeting report, 8–10 March 2023. 2023.

18. Korgaonkar NS, Kumar A, Yadav RS, Kabadi D, Dash AP. Mosquito biting activity on humans & detection of Plasmodium falciparum infection in Anopheles stephensi in Goa, India. Indian J Med Res. 2012 Jan;135(1):120–6.

19. Shahandeh K, Basseri H, Pakari A, Riazi A. Mosquito Vector Biting and Community Protection in a Malarious Area, Siahoo District, Hormozgan, Iran. Iran J Arthropod-Borne Dis. 2010 Dec 31;4(2):35–41.

20. Coetzee M. Key to the females of Afrotropical Anopheles mosquitoes (Diptera: Culicidae). Malar J. 2020 Feb 13;19(1):70.

21. Rund SSC, Bonar NA, Champion MM, Ghazi JP, Houk CM, Leming MT, et al. Daily rhythms in antennal protein and olfactory sensitivity in the malaria mosquito Anopheles gambiae. Sci Rep. 2013 Aug 29;3(1):2494.

22. Rund SSC, Lee SJ, Bush BR, Duffield GE. Strain- and sex-specific differences in daily flight activity and the circadian clock of *Anopheles gambiae* mosquitoes. J Insect Physiol. 2012 Dec 1;58(12):1609–19.

23. Petrić D, Bellini R, Scholte EJ, Rakotoarivony LM, Schaffner F. Monitoring population and environmental parameters of invasive mosquito species in Europe. Parasit Vectors. 2014 Apr 16;7(1):187.

24. Parker JD, Torchin ME, Hufbauer RA, Lemoine NP, Alba C, Blumenthal DM, et al. Do invasive species perform better in their new ranges? Ecology. 2013 May;94(5):985–94.

25. Wilson AL, Courtenay O, Kelly-Hope LA, Scott TW, Takken W, Torr SJ, et al. The importance of vector control for the control and elimination of vector-borne diseases. PLoS Negl Trop Dis. 2020 Jan 16;14(1):e0007831.

26. Sangbakembi-Ngounou C, Costantini C, Longo-Pendy NM, Ngoagouni C, Akone-Ella O, Rahola N, et al. Diurnal biting of malaria mosquitoes in the Central African Republic indicates residual transmission may be “out of control.” Proc Natl Acad Sci U S A. 2022 May 24;119(21):e2104282119.

27. Duffield GE. Circadian and daily rhythms of disease vector mosquitoes. Curr Opin Insect Sci. 2024 June 1;63:101179.

28. Das S, Dimopoulos G. Molecular analysis of photic inhibition of blood-feeding in Anopheles gambiae. BMC Physiol. 2008 Dec 16;8:23.

29. Degefa T, Githeko AK, Lee MC, Yan G, Yewhalaw D. Patterns of human exposure to early evening and outdoor biting mosquitoes and residual malaria transmission in Ethiopia. Acta Trop. 2021 Apr;216:105837.

30. Esayas E, Gowelo S, Assefa M, Vajda EA, Thomsen E, Getachew A, et al. Impact of nighttime human behavior on exposure to malaria vectors and effectiveness of using long-lasting insecticidal nets in the Ethiopian lowlands and highlands. Parasit Vectors. 2024 Dec 18;17(1):520.

31. Taylor R, Messenger LA, Abeku TA, Clarke SE, Yadav RS, Lines J. Invasive *Anopheles stephensi* in Africa: insights from Asia. Trends Parasitol. 2024 Aug 1;40(8):731–43.

32. Ng’habi KR, Lee Y, Knols BGJ, Mwasheshi D, Lanzaro GC, Ferguson HM. Colonization of malaria vectors under semi-field conditions as a strategy for maintaining genetic and phenotypic similarity with wild populations. Malar J. 2015 Jan 21;14:10.

33. Baeshen R, Ekechukwu NE, Toure M, Paton D, Coulibaly M, Traoré SF, et al. Differential effects of inbreeding and selection on male reproductive phenotype associated with the colonization and laboratory maintenance of Anopheles gambiae. Malar J. 2014 Jan 13;13(1):19.

34. Bedasso AH, Gutto AA, Waldetensai A, Eukubay A, Bokore GE, Kinde S, et al. Malaria vector feeding, peak biting time and resting place preference behaviors in line with Indoor based intervention tools and its implication: scenario from selected sentinel sites of Ethiopia. Heliyon. 2022 Dec 1;8(12):e12178.

35. Odero JI, Abong’o B, Moshi V, Ekodir S, Harvey SA, Ochomo E, et al. Early morning anopheline mosquito biting, a potential driver of malaria transmission in Busia County, western Kenya. Malar J. 2024 Mar 4;23(1):66.

36. Nzioki I, Machani MG, Onyango SA, Kabui KK, Githeko AK, Ochomo E, et al. Differences in malaria vector biting behavior and changing vulnerability to malaria transmission in contrasting ecosystems of western Kenya. Parasit Vectors. 2023 Oct 21;16(1):376.

37. Govella NJ, Ferguson H. Why Use of Interventions Targeting Outdoor Biting Mosquitoes will be Necessary to Achieve Malaria Elimination. Front Physiol. 2012 June 12;3:199.

38. Killeen GF, Chitnis N. Potential causes and consequences of behavioural resilience and resistance in malaria vector populations: a mathematical modelling analysis. Malar J. 2014 Mar 14;13(1):97.

39. Cooke MK, Kahindi SC, Oriango RM, Owaga C, Ayoma E, Mabuka D, et al. ‘A bite before bed’: exposure to malaria vectors outside the times of net use in the highlands of western Kenya. Malar J. 2015 June 25;14(1):259.

40. Tadesse FG, Ashine T, Teka H, Esayas E, Messenger LA, Chali W, et al. Anopheles stephensi Mosquitoes as Vectors of Plasmodium vivax and falciparum, Horn of Africa, 2019. Emerg Infect Dis. 2021 Feb;27(2):603–7.

41. Emiru T, Getachew D, Murphy M, Sedda L, Ejigu LA, Bulto MG, et al. Evidence for a role of Anopheles stephensi in the spread of drug- and diagnosis-resistant malaria in Africa. Nat Med. 2023 Dec;29(12):3203–11.

42. Zhou G, Zhong D, Yewhalaw D, Yan G. Anopheles stephensi ecology and control in Africa. Trends Parasitol. 2024 Feb;40(2):102–5.

43. Ochomo EO, Gimnig JE, Awori Q, Abong’o B, Oria P, Ashitiba NK, et al. Effect of a spatial repellent on malaria incidence in an area of western Kenya characterised by high malaria transmission, insecticide resistance, and universal coverage of insecticide treated nets (part of the AEGIS Consortium): a cluster-randomised, controlled trial. Lancet Lond Engl. 2025 Jan 11;405(10473):147–56.

44. Knols BG, Njiru BN, Mathenge EM, Mukabana WR, Beier JC, Killeen GF. MalariaSphere: A greenhouse-enclosed simulation of a natural Anopheles gambiae (Diptera: Culicidae) ecosystem in western Kenya. Malar J. 2002 Dec 18;1(1):19.

45. Ferguson HM, Ng’habi KR, Walder T, Kadungula D, Moore SJ, Lyimo I, et al. Establishment of a large semi-field system for experimental study of African malaria vector ecology and control in Tanzania. Malar J. 2008 Aug 20;7(1):158.

